# Sharpened visual memory representations are reflected in inferotemporal cortex

**DOI:** 10.1101/2025.04.28.651105

**Authors:** Barnes G.L. Jannuzi, Catrina M. Hacker, Simon Bohn, Travis Meyer, Madison L. Hay, Nicole C. Rust

## Abstract

Humans and other primates can robustly report whether they’ve seen specific images before, even when those images are extremely similar to ones they’ve previously seen. Multiple lines of evidence suggest that pattern separation computations in the hippocampus (HC) contribute to this behavior by shaping the fidelity of visual memory. However, unclear is whether HC uniquely determines memory fidelity or whether computations in other brain areas also contribute. To investigate, we recorded neural signals from inferotemporal cortex (ITC) and HC of two rhesus monkeys as they performed a memory task in which they judged whether images were novel or exactly repeated in the presence of visually similar lure images with a range of visual similarities. We found behavioral evidence for sharpening, reflected as memory performance that was nonlinearly transformed relative to a benchmark defined by visual representations in ITC. As expected, we found that behavioral sharpening aligned with visual memory representations in HC. Surprisingly, and unaccounted for by HC pattern separation proposals, we also found neural correlates of behavioral sharpening reflected in ITC. These results, coupled with further analysis of the data, suggest that ITC contributes to shaping the fidelity of visual memory in the transformation from visual processing to memory storage and signaling.

**Significance:** Visual recognition memories are stored with remarkable visual fidelity, allowing humans and other primates to distinguish images they have encountered from visually similar images they have not. This fidelity has long been attributed to computations in the hippocampus that sharpen visual representations before memory storage (“pattern separation”). Unclear is how this proposal aligns with other evidence that visual memories are stored within high-level visual cortex itself, before signals reach the hippocampus. Here we demonstrate that, like the hippocampus, inferotemporal cortex also reflects sharpened visual memory representations, suggesting that visual cortex contributes to shaping the visual fidelity of visual memory.

## Introduction

Humans (and other primates) have a remarkable ability to remember the images they have seen. Subjects can correctly report whether they’ve seen specific images after seeing thousands of them, each only once and for a few seconds (Standing, 1973; Brady et al., 2008). In addition, subjects typically remember images with impressive visual detail, even in the absence of specific instructions to do so (Standing, 1973; Vogt and Magnussen, 2007; Brady et al., 2008; Konkle et al., 2010; Stark et al., 2015, 2019). How do our brains manage to support the astounding visual specificity of visual recognition memory? (Below, we refer to visual recognition memory as “visual memory” for ease of reading.)

Several lines of evidence suggest a prominent role of the hippocampus (HC) in shaping the fidelity of visual memory. When visual memory is tested in patients with HC damage, they have preserved memory for visually distinct images, but they falsely remember lure images that are similar to those already seen at higher rates than controls (Stark et al., 2013, 2015, 2019).

Similarly, individuals with an array of HC-implicated disorders display impaired memory performance for similar images, including patients with age-related mild cognitive impairment (Stark et al., 2013; Bennett et al., 2019) and Alzheimer’s disease (Ally et al., 2013; Monti et al., 2014). Likewise, fMRI comparisons of visual representations between the HC and a primary input, entorhinal cortex, suggest that hippocampal memory representations are more sparse, implying computations within the hippocampus sharpen visual representations (Bakker et al., 2008; Lacy et al., 2011; Berron et al., 2016).

Taken together, these results lead to the now-popular “pattern separation hypothesis” for how HC may shape visual memory by sparsifying visual representations. The idea begins with the notion that visual representations of similar images in high-level visual cortex (including inferotemporal cortex (ITC)) are highly overlapping (Figure 1, left), and because memory storage and signaling is a noisy, information-degrading process, these representations cannot support the visual specificity of visual memory (Yassa and Stark, 2011; Leal and Yassa, 2018; Stark et al., 2019). To resolve this downstream of ITC, pattern separation computations in the HC subregion dentate gyrus transform overlapping and distributed representations arriving from entorhinal cortex into more sparsely distributed representations in HC subregion CA3 (Figure 1, right). These dissimilar visual representations are then protected from confusion during noisy memory storage and signaling. Consistent with this proposal, one study found that visual memory behavior in monkeys (measured by preferential looking for novel as compared to repeated stimuli) could be decoded from CA3 (Sakon and Suzuki, 2019).

**Figure 1.**
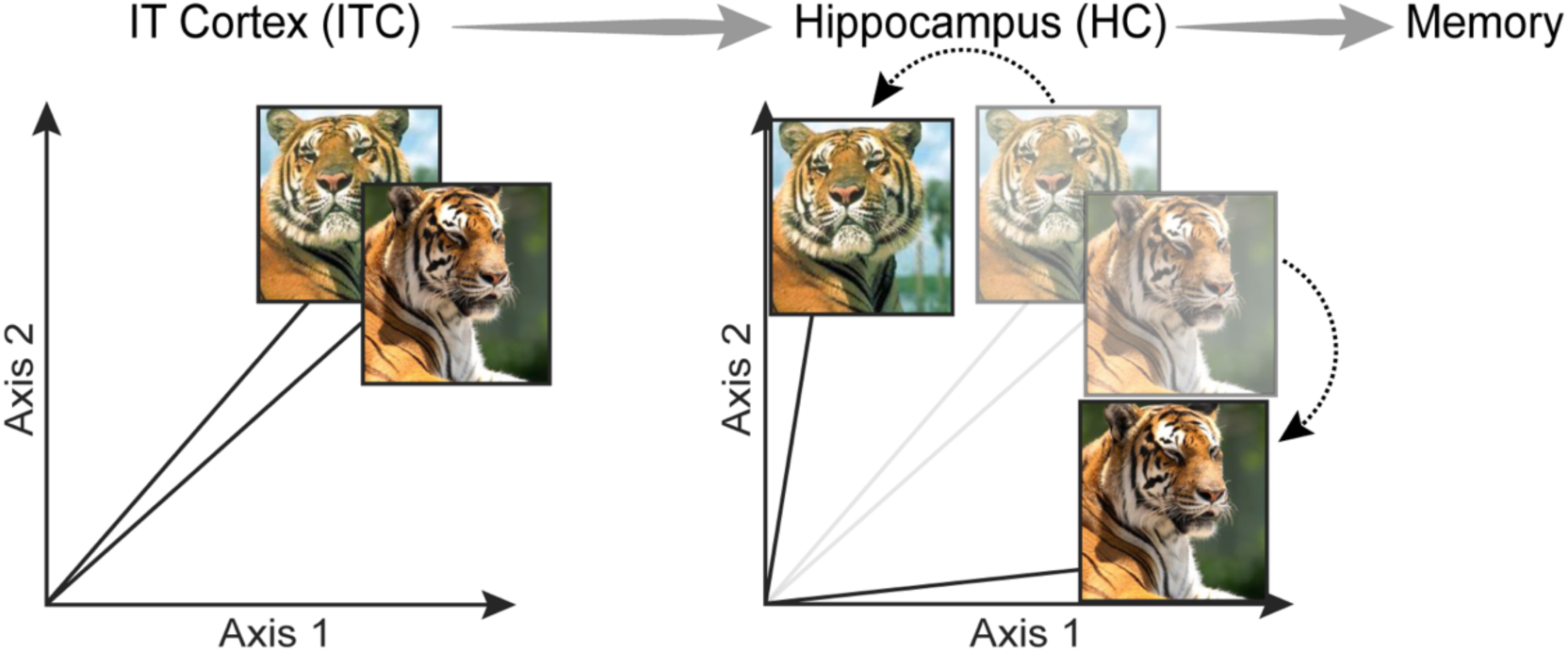
The hippocampal pattern separation hypothesis. Left: Hypothetical ITC population responses to two tiger images that are highly overlapping. Because of their high similarity, they have highly overlapping neural representations. The visual representations of these two images exist in high-d population space (dimensionality equal to the number of ITC neurons), but only two dimensions are visualized here. Because memory storage and signaling is a noisy process, confusion can follow from highly similar images like these, where an individual falsely reports seeing one image when in truth, they saw the other. *Right:* Hypothetical population responses to the same two images in HC following pattern separation. Because the images are less overlapping, memory is less susceptible to noise. Dotted arrows emphasize the transformation in the population representations between ITC and HC.

While evidence implicating HC in supporting visual memory fidelity is compelling, it seems to contradict evidence that visual memories are stored within high-level visual cortex itself, before signals reach HC. Namely, a number of studies have demonstrated an alignment between visual memory behavior and repetition suppression (RS), an adaptation-like response reduction that occurs in ITC upon repeating a stimulus (Fahy et al., 1993; Li et al., 1993; Miller and Desimone, 1994; Xiang and Brown, 1998; Meyer and Rust, 2018; Mehrpour et al., 2021), thereby supporting ITC RS as a candidate signal that the brain uses to drive visual memory. It is possible that fidelity-shaping computations within ITC also contribute to the transformation from vision to visual memory. However, because ITC visual memory representations of highly similar images have never been systematically compared with the fidelity of visual memory behavior, this remains unclear.

Here, we ask whether and how transformations between ITC visual representations and HC visual memory representations occur. To achieve this, we trained monkeys to perform a visual memory task in which they reported familiarity for novel and exactly repeated images, as well as visually similar lures (which were benchmarked for their visual similarity against ITC visual representations). As monkeys performed this task, we recorded neural data from HC and ITC. We found robust behavioral evidence of sharpening, reflected as nonlinearly transformed memory performance on lure images relative to a model of ITC visual representations (Kubilius et al., 2019). As expected from previous work, we found that visual memory representations in HC aligned with visual memory behavior. Surprisingly, we also found evidence of sharpening in ITC. These results, coupled with additional analyses, suggest that ITC contributes to shaping the fidelity of visual memory in the transformation from visual to memory representations.

## Results

### The adapted mnemonic similarity task

The monkeys’ behavioral task was an adaptation of the Mnemonic Similarity Task (MST) that has been used extensively to measure the contribution of HC to shaping the fidelity of visual memory (Kirwan et al., 2007; Bakker et al., 2008). In the original task, subjects viewed images that were novel, exactly repeated, or similar to those viewed before, where degree of similarity was determined empirically as the likelihood that a lure image was mistakenly classified as repeated, on average, across many subjects. Here, we take a complementary approach where we measure similarity by benchmarking against representational dissimilarity based on a state-of-the-art model of ITC: CORnet_S-IT (Kubilius et al., 2019)(see Methods). This network has been shown to be predictive of neural data from macaques observing images, and behavioral predictions on object identification tasks from human participants (Kubilius et al., 2019). The rationale behind relying on a model of ITC for these experiments is two-fold. First, our single-exposure memory experiments require exceedingly large numbers of lure pair images (∼30,000) at a range of visual dissimilarities, which is a prohibitively large number to test and curate empirically. Second, benchmarking against ITC representations allows us to assess from behavior alone, the extent, if at all, that nonlinear sharpening computations happen downstream of ITC visual representations.

In this task, monkeys viewed images and reported whether they were novel (i.e., never before seen in training or testing) or repeated (meaning that they had seen that image exactly once before and earlier in the experimental session) (Figure 2a-b). Monkeys initiated each trial by fixating a point at the center of the screen, followed by a brief delay and then the presentation of an image (Figure 2a). After 500ms of fixating on the image, a go cue appeared, indicating that the monkeys were free to make their selection via a saccade to one of two response targets.

**Figure 2.**
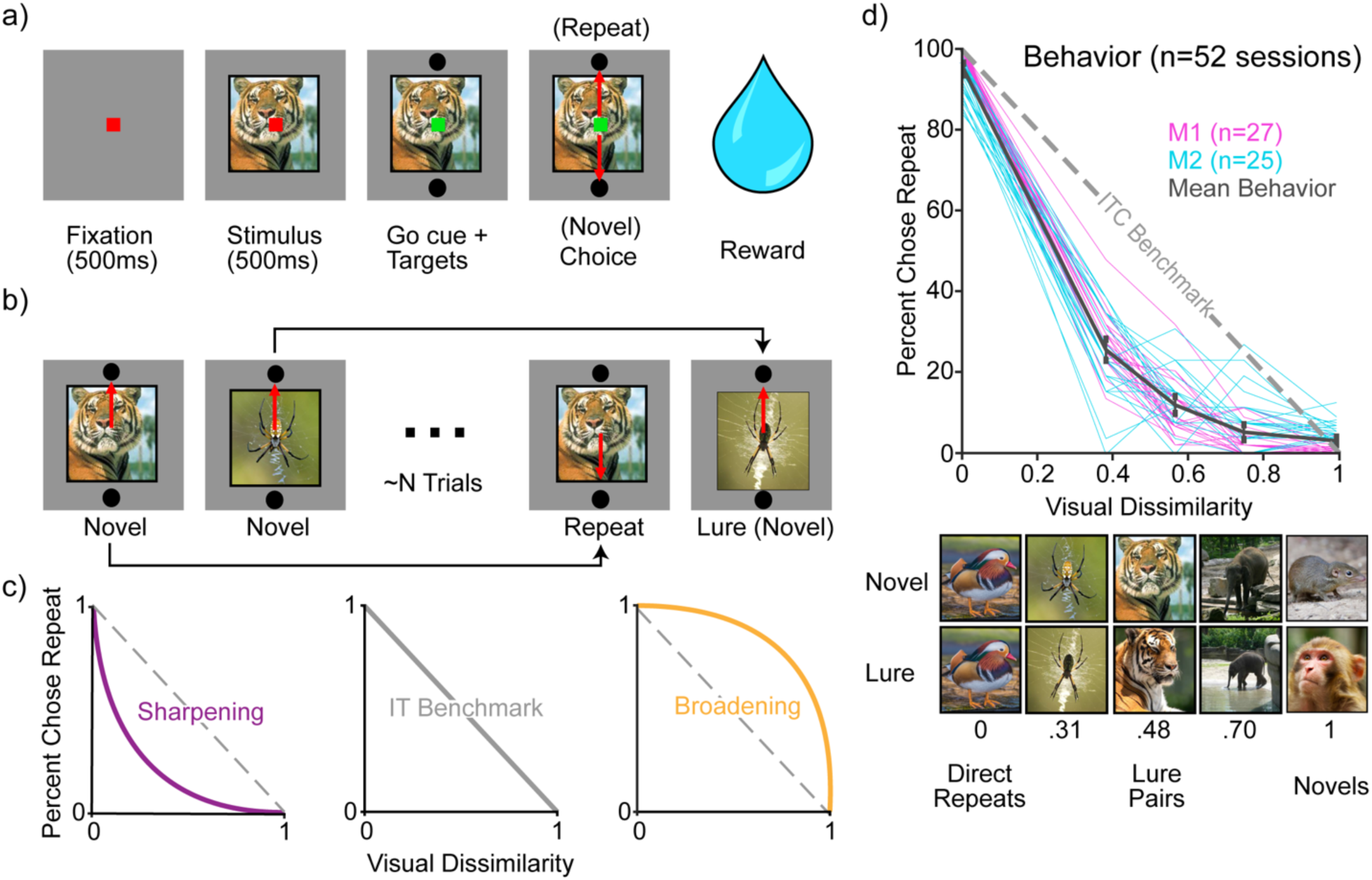
Adapted Mnemonic Similarity Task (MST) **a)**. Each trial began with the monkey fixating a red dot for 500ms. If fixation was successfully maintained, an image was presented for 500ms before the fixation square turned green and targets appeared above and below the image. The monkey was then free to make a saccade to one of the two targets to report whether they judged the image to be novel or repeated. Successful choices were rewarded with juice. **b)** Task overview: An example sequence of trials which include novel, exactly repeated, and lure images were separated by N trials where N was adjusted to equate performance between the monkeys (N= ∼24 trials in monkey 1 and ∼8 trials in monkey 2). Black arrows connect novel/repeat and novel/lure pairs. **c)** Hypothetical behavioral patterns illustrating different possible outcomes. The diagonal gray dashed line (middle) represents the expected performance of the monkey if they performed exactly as well as would be expected based on our measure of ITC dissimilarity. Shown are hypothetical examples of behavioral sharpening (left) and broadening (right) relative to this benchmark. **d)** Behavioral performance. *Top:* Monkey 1 and Monkey 2 sessions plotted in magenta and cyan, respectively. Mean behavioral performance across all 52 sessions is plotted in black with 95% confidence interval error bars. Behavioral reports of the monkeys indicate evidence of sharpening on average and within individual sessions. *Bottom:* example lure image pairs at a range of visual dissimilarities. Dissimilarity values were quantified by the similarity of lure pairs based on a model of ITC: CORnet_S-IT(see Methods).

Correct responses were rewarded with juice. Lure images were similar but not identical to novel images and contained an object from the same category, systematically titrated at a range of visual similarities (Figure 2d, bottom). Only a small fraction (13%) of images were lures.

The first presentation of a lure image was rewarded when the monkey chose “novel.” Lure images were also repeated, and on those trials, they were rewarded when the monkey chose “repeat”. (Non-lure) repeated images, lure images, and lure repeats were shown ∼N trials following their lure pair, where N was adjusted to equate the performance across the two monkeys (∼24 trials for monkey 1 and ∼8 trials for monkey 2). Novel images that were paired with lures were never repeated. While the first image presented in each session was always novel, the probability of subsequent images being novel versus repeated quickly converged to 57%. The slight increase in novel images is because lure images are, by definition, novel; this bias was mitigated by repeating the lure images, however, the first novel image in a lure pair was not repeated. In each behavioral session, monkeys viewed 1500 images consisting of novel, exact repeats, lures, and lure repeats.

Performance was measured as the percent chose repeat (PCR) for any given trial category (Figure 2c-d). For novel images, this represents the false alarm rate, and for repeated images, this represents the hit rate. For lure images, the PCR is the rate of false memory, or equivalently the mnemonic confusability of the lure images.

Assessing hypotheses about “sharpening” (or broadening) downstream of ITC visual representations requires specifying a benchmark of “no sharpening” to measure against. Here we use a method inspired by representational similarity analysis (Kriegeskorte et al., 2008) where we measured dissimilarity based on ITC visual representations. Specifically, we used: 1 - the Pearson’s correlation of the ITC population responses on two trials, where the ITC responses are either estimated by an ITC model or are measured directly or indirectly from brain activity (Kriegeskorte et al., 2008) (see Methods). This can be thought of as plotting visual representations on the x-axis and memory representations on the y-axis. If visual memory behavior aligns with this benchmark, it indicates that the transformation from ITC visual representations to visual memory behavior is linear. Conversely, deviations from this benchmark indicate that this transformation is shaped by nonlinear computations (which may include hippocampal pattern separation).

We found clear and robust evidence for behavioral sharpening for both monkeys, including within individual sessions (Figure 2d; magenta and cyan lines). This sharpening could follow in whole or in part from hippocampal pattern separation computations, or it could follow from additional computations at one or more earlier stages in the processing hierarchy (which includes computations within ITC itself, as well as perirhinal cortex, parahippocampal cortex, and entorhinal cortex) (Felleman and Van Essen, 1991). To investigate, we compared sharpening behavior with memory representations at the ends of this processing chain: in HC and ITC.

### Neural analogs of sharpening in HC and ITC

As the monkeys performed the task, we recorded neural spiking responses from HC or ITC using a multi-channel electrode acutely lowered before the monkey began each session. For quality control, each recording session was screened based on its recording stability across the session, the number of behavioral trials completed, visual responsiveness, and memory information (see Methods). Data reported here correspond to the subset of images that have a valid behavioral response — meaning the monkey completed the trial without breaking fixation — for both the first and second presentations of the image (either a novel/exact repeated or a novel/lure pair).

Comparisons between neural responses and behavior in single-trial memory tasks like this require many hundreds of units (Meyer and Rust, 2018). To analyze the neural data, we concatenated units across sessions to create larger pseudopopulations. When creating these pseudopopulations, we aligned novel, repeated and lure images based on their lure dissimilarity (see Methods). The resulting pseudopopulations consisted of the responses to novel and repeated images, as well as pairs of novel and lure images. These populations were matched for their numbers of image pairs across brain regions within each monkey and condition (resulting in 213 and 142 novel/repeated image pairs and 133 and 80 novel/lure image pairs in monkey 1 and monkey 2 respectively). Within each monkey and across the two brain areas, the number of units included in each population was matched for memory information using classifier performance (see Methods). The resulting dataset included n=422 HC and n=504 ITC units for monkey 1 and n=340 HC units and n=595 ITC units for monkey 2.

Having finalized the neural populations, we next sought to relate them directly to behavior. A decoding scheme is required to relate neural representations of memory with memory behavior. When memory is reflected as RS, that decoding scheme can be approximated by a simple total spike count linear decoder or ’spike count classifier’ (SCC); however, when memory is reflected as mixtures of RS in some units and repetition enhancement (RE) in others, that linear decoder requires mixtures of positive and negative weights. Following on earlier work suggesting that visual memory information is reflected in ITC as RS, we hypothesized that this is true in HC as well.

An examination of the grand mean firing rate responses to novel and repeated images (Figure 3a), revealed that RS increased in strength over the 500ms. We thus chose to evaluate memory performance by counting spikes in a window 300-500ms following stimulus onset (and unless stated otherwise, this holds for all analyses presented). For each unit, d’ was calculated by comparing the novel and repeated responses (see Methods). We found that these distributions of d’ were roughly normal but shifted towards positive values in both HC and ITC, indicating populations dominated by RS along with some noise (Figure 3b-c).

**Figure 3.**
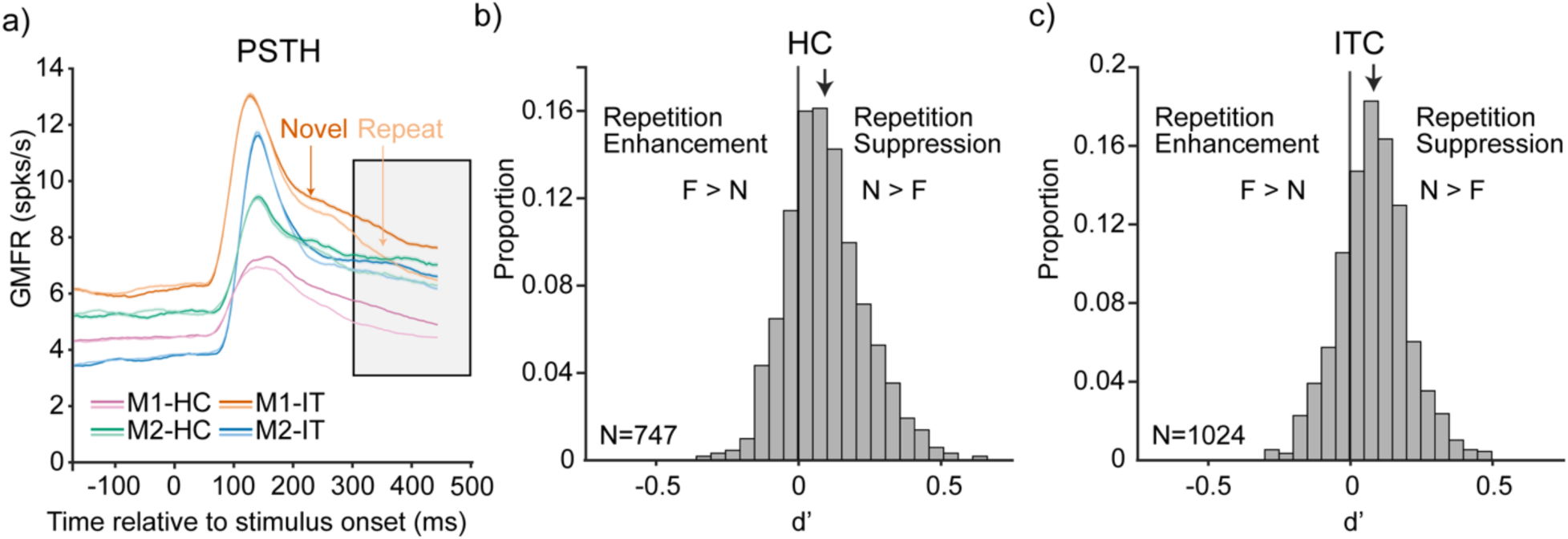
Memory information is reflected as RS in both HC and ITC. **a)** Grand mean peristimulus time histogram (PSTH) of neural responses to novel and repeated images, averaged across all units in each population and response type. Traces smoothed by averaging over a 50ms time bin. Monkey1-HC, Monkey2-HC, Monkey1-IT, Monkey2-IT, are plotted as Magenta, Green, Orange, and Blue respectively. Novel images are plotted as darker lines and repeated images are plotted as lighter lines. Standard error of the means plotted as tight areas around lines. Gray shaded box (300ms-500ms) indicates the time window over which we computed d’. **b-c)** Histograms of d’ values for individual units in HC and ITC. Arrows indicate population means (HC = 0.090; ITC = 0.086). Values greater than zero have positive d’, indicating RS. Values less than zero have negative d’, indicating RE.

To determine whether any of the negative d’ units reflected reliable memory information as RE (as opposed to noise), we compared cross-validated classifier performance for a weighted linear decoder that can take the sign of each unit into account by assigning RE and RS units opposite signs. Specifically, we computed memory classification performance as we added units ranked by their signed classifier weights as: high-to-low (red), low-to-high (blue) or randomly selected (black), based on training data alone (Figure 4). In these plots, above-chance performance for small numbers of units with low-to-high ranking would be indicative of RE contributing to classifier performance, whereas above-chance performance for small numbers of units with high-to-low ranking is indicative of RS contributing to classifier performance. We found that trends in performance were dominated by the latter pattern, with negligible evidence for the former in HC and ITC of both monkeys (Figure 4). Together, these results suggest that memory information in both HC and ITC is reflected as RS (as opposed to mixtures of RS in some units and RE in others). Below, we compare neural memory representations with behavior using a total spike count linear classifier (which decodes RS).

**Figure 4.**
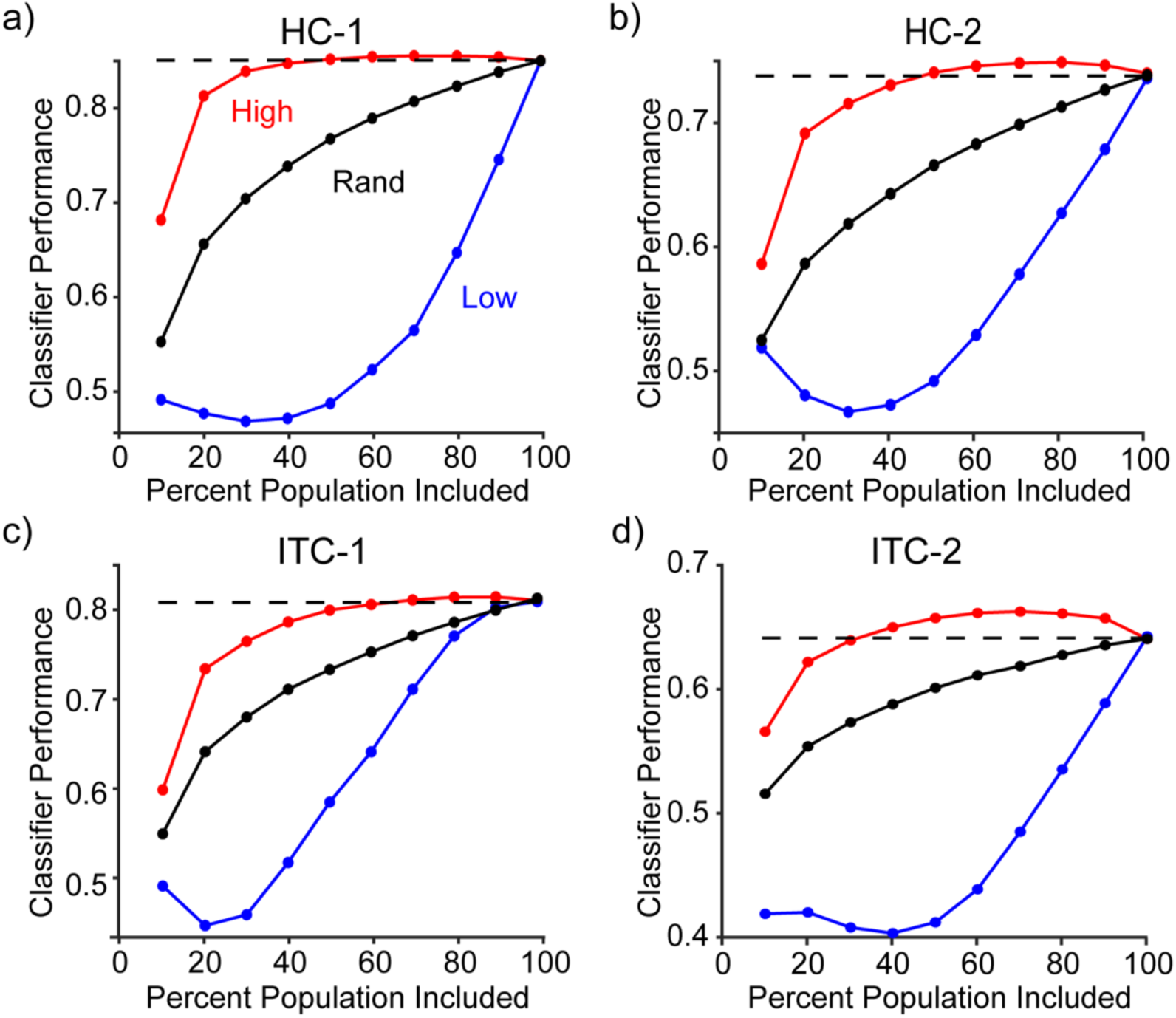
Population-based confirmation that memory information is reflected in HC and ITC as repetition suppression. **a-d)** Performance of a weighted linear classifier as a function of population size with respect to units ranked by classifier weight. Each plot shows performance as units are added to the population that are ranked by signed classifier weights high-to-low (‘High’-red; where first-ranks are RS), low-to-high (‘Low’-blue; where first-ranks are RE), or randomly (‘Rand’-black). Full-sized population classifier performance is reflected by the dashed horizontal line. **a-b)** Hippocampus populations for monkey 1 (left) and monkey 2 (right). **c-d)** The same, but for ITC populations.

To compare neural representations with behavior, the spike count decoder was trained on novel and repeated images and performance was assessed both with cross-validated images from the same conditions as well as for its ability to generalize to lure conditions. We found that classifier predictions based on HC neural responses aligned with behavioral sharpening in both animals (Figure 5a-b). More surprising was that classifier predictions based on ITC neural responses *also* clearly reflected robust sharpening relative to the benchmark of ITC visual representations in both monkeys (Figure 5c-d). Importantly, that the neural predictions correspond with the sharpened behavior (rather than the benchmark predicted by ITC visual representations) suggests the existence of a nonlinear transformation between ITC visual and memory representations.

**Figure 5.**
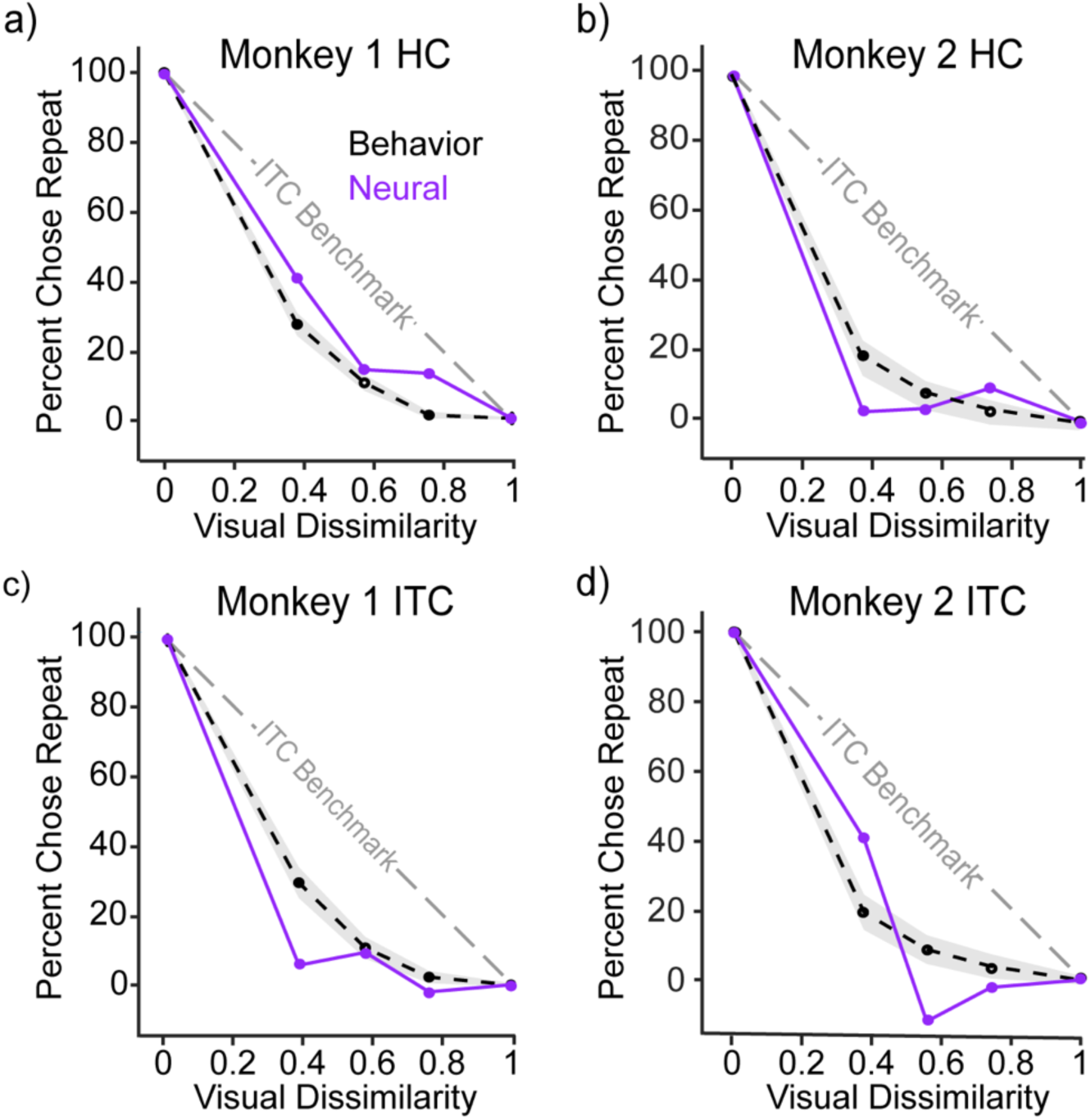
Behavioral sharpening is reflected in both HC and ITC neural representations of memory. **a-d)** The comparison of sharpening behavior (black, dashed), cross-validated spike count classifier performance (purple), and ITC benchmark (gray)The black dashed line shows the mean behavioral performance across all sessions with 95% confidence intervals shown with a gray shaded region. The purple line reflects classifier predictions of behavior, classifier predictions have been rescaled such that novel and repeated performance is matched to behavior and lure prediction is scaled proportionally (see Methods).

To validate that these results were not due to individual units with large sporadic firing rates, we re-ran the analysis after normalizing firing rates for each unit by z-scoring the spike firing rate responses across all images. This manipulation did not qualitatively change the results (Supplemental Figure 1). As expected from the nature of spike count decoder, classifier performance aligned with the distributions of spike firing rates for each of the lure distributions (Supplemental Figure 2).

Finally, we wondered whether HC and ITC visual memory representations emerged relative to stimulus onset as sharpened (as opposed to linearly with sharpening later) and whether there were any obvious differences in the dynamics of how they emerged in HC versus ITC. For these analyses, we combined the data from the two monkeys into one population to maximize SNR and the sensitivity to detect differences. We measured sharpening based on a metric that computed area of the neural classifier prediction curve relative to the ITC benchmark, weighted by overall classifier performance (see Methods). With this metric, positive values reflect sharpening whereas negative values reflect broadening (Figure 2c). Consistent with the intuitions presented in Figure 5, this metric reflected sharpening in both HC and ITC at the end of the viewing window that approximately aligned with behavior (Figure 6a, gray arrow at 400 ms, reflecting a spike count window 300-500 ms). Likewise, visual memory representations in HC and ITC emerged with approximately the same dynamics (Figure 6a, arrows at 150, 275, 350 and 400 ms; Figure 6b). Notably, we found no clear evidence of sharpening emerging first in HC, followed by ITC, suggesting that the sharpening observed in ITC is not just a result of responses sharpened in HC that are fed back to ITC.

**Figure 6.**
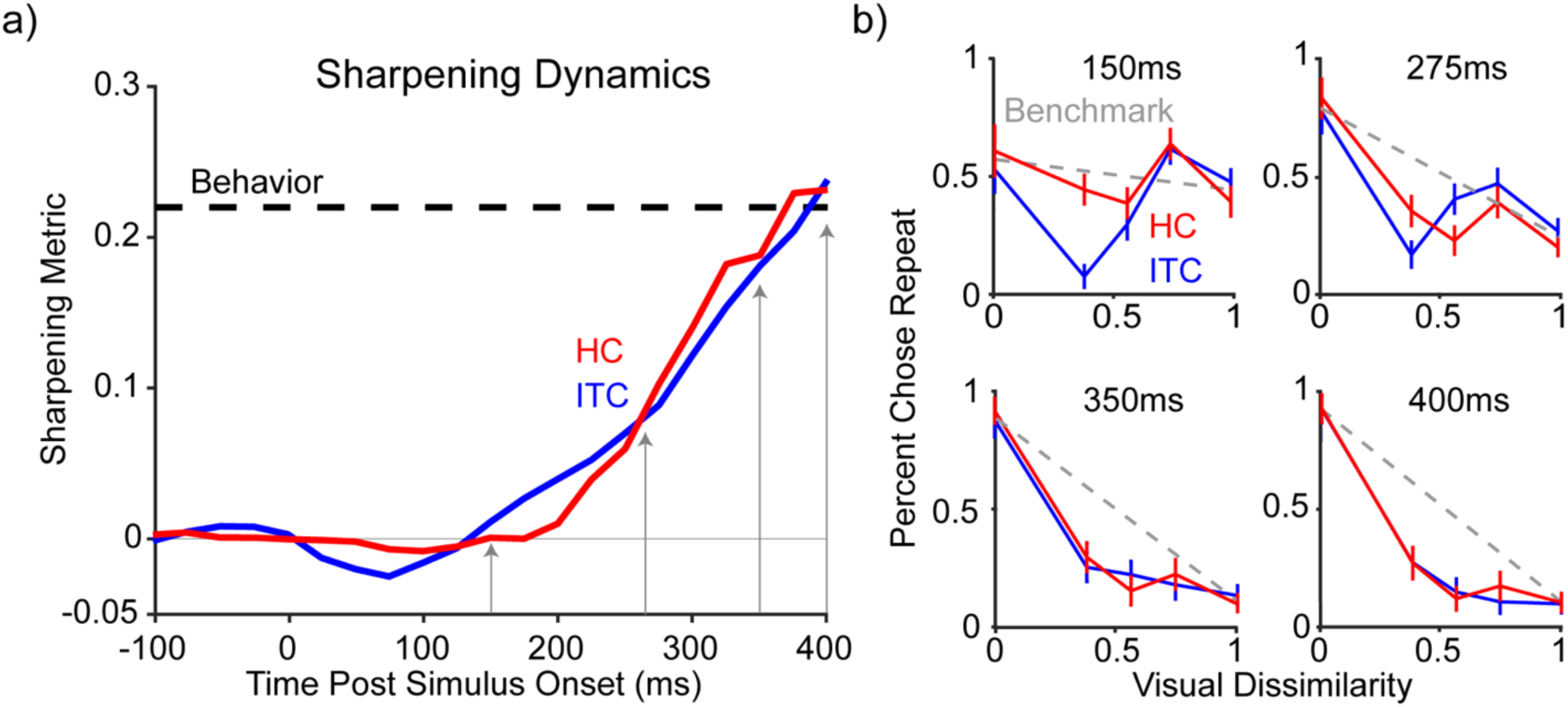
Neural sharpening emerges in both HC and ITC. **a-b)** Dynamics analyses performed using a 200ms window on a combined population of both monkeys across HC and ITC (Red and Blue respectively). **a)** Dynamics of sharpened classifier performance (see methods for sharpening metric). Behavioral sharpening plotted as a black dashed line. Gray arrows indicate centers of time slices shown in panel b. **b)** Individual 200ms time slices of classifier accuracy as a function of dissimilarity. The dashed gray line connects average HC and IT performance for novel and repeated images, or equivalently, the ITC benchmark. To illustrate classifier performance to novel and repeat images during each slice, classifier performances (and thus, the ITC benchmark) have not been rescaled here. Error bars indicate standard error of the mean over classifier cross-validation iterations.

All of these results rely on comparison with the benchmark of ITC visual representations generated using CORnet_S-IT as a model (Kubilius et al., 2019). However, there are limitations of how effective artificial neural networks serve as models for brain region representations (Schaeffer et al., 2022; Ahlert et al., 2024; Feghhi et al., 2024), and of particular concern for this experiment are model inaccuracies that nonlinearly reflect actual ITC representational similarity, as these could spuriously lead to the conclusion that visual memory representations are sharpened. Indeed, if a nonlinear relationship exists between CORnet_S-IT dissimilarity and ground-truth neural dissimilarity, the sharpening we’ve observed (Figure 2,5,6) could be an artifact. While our experiment was not optimized to estimate neural similarity (as each image was presented at most twice), the data do have some power to make these estimates. To test for nonlinearities, we thus compared CORnet_S-IT and neural estimates of dissimilarity for novel-lure pairs, which were well-described as linear with no indication of a nonlinear relationship (Supplemental Figure 3).

## Discussion

This study aimed to test the nature of the transformations from ITC visual representations to HC visual memory representations. Based on prior work (Yassa and Stark, 2011; Bennett et al., 2019; Sakon and Suzuki, 2019), we expected to find that HC visual memory representations mimicked behavioral sharpening, and indeed, we found evidence for this in both monkeys.

However, an open question is the extent that hippocampal pattern separation was primarily responsible for this nonlinear transformation, or if other brain regions such as ITC also reflect some degree of sharpening — surprisingly, we found this to clearly be the case.

Having found robust evidence for sharpening in ITC visual memory representations, reflected as RS, the central question is: What might the source of sharpened ITC RS be? Here, there are four possibilities to consider. The first two are related to processing before or within ITC. First, ITC RS might be the result of cumulative RS computations in the ventral visual pathway prior to ITC (from structures such as V2 and V4), which have been shown to be independent of ITC RS (Williams and Olson, 2022). However, RS in the V2 input to ITC is weaker than within ITC itself, and thus cannot alone account for it (Williams and Olson, 2022). A second possibility is that ITC RS is the result of nonlinear computations within ITC itself (Brown et al., 1987). RS is inherently an adaptation-like phenomenon that extends over long timescales, and its alignment with familiarity behavior is captured with models in which familiarity memory happens via feedforward, anti-Hebbian plasticity in an IT-like network layer (Földiák, 1990; Tyulmankov et al., 2022). Likewise, the dynamics of RS are captured by models in which familiarity memories are stored as changes in ITC synaptic weights, and RS emerges through recurrent interactions between excitatory and inhibitory neurons (Lim et al., 2015; Agnes and Vogels, 2024). In this proposal, sharpened RS emerges as part of this ITC computation.

The third possibility is that sharpened ITC visual memory representations reflect computations in the medial temporal lobe that are then fed back onto ITC. This could be the result of RS computations in structures such as perirhinal cortex (Bogacz and Brown, 2003), where evidence of feedback to ITC exists for other forms of memory (paired associations, (Miyashita, 2019)).

Likewise, memory representations in ITC could reflect computations that happen in HC that fed back to cortex, analogous to how other forms of learning are thought to occur (Kim et al., 2023). A final possibility is that sharpened ITC visual memory representations follow from computations that rely on reciprocal interactions between multiple brain areas that include ITC as well as one or more structures such as perirhinal cortex, entorhinal cortex, HC, and/or prefrontal cortex. (Ranganath, 2006; Newmark et al., 2013; Li et al., 2015; Sučević and Schapiro, 2023). From our data alone, we are unable to fully clarify the contributions from each class of model towards sharpened RS in ITC. However, a parsimonious explanation of our results is that the existence of sharpened RS in ITC is not merely an epiphenomenon (and a metabolically wasteful feedback step in the brain) but is indicative of a contribution of ITC to shaping the fidelity of visual memory. Future experiments will be required to tease apart exactly what that contribution is.

A significant body of research in humans suggests that an intact and functioning HC is required to support the fidelity of visual memory (Kirwan et al., 2007; Bakker et al., 2008; Wixted and Squire, 2010; Lacy et al., 2011; Yassa and Stark, 2011; Stark et al., 2013, 2019; Suthana et al., 2015; Barbeau et al., 2017; Bennett et al., 2019). Namely, individuals with impairments to HC are more likely to mistakenly report that they’ve seen visually-similar lures than control subjects. Likewise, evidence from human fMRI (Nash et al., 2021) and single unit recordings (Suthana et al., 2015) suggest that visual memory representations are more sharpened in HC than its input, entorhinal cortex. One mystery of our results is why we fail to qualitatively replicate this result, as visual memory representations were no less sharpened in ITC than HC in our data. One possible explanation is a species difference — although evidence suggests that “pattern separation” computations happen even in rodents in the context of spatial recognition tasks (van Goethem et al., 2018), and that rodent HC is required for innate object recognition memory behavior (Broadbent et al., 2010; Cohen et al., 2013; Cohen and Stackman, 2015; Johnson et al., 2017). Another possible explanation is that HC may contribute to sharpening when conceptual (rather than just perceptual) factors are at play (Barense et al., 2007; Bonnen et al., 2021; Gurguryan et al., 2024), acknowledging that the monkeys in this study were largely unfamiliar with the objects presented in the images used in these experiments (such as firetrucks and giraffes). Of note is that neither of these possible scenarios undermines the conclusion that ITC contributes in some way to shaping the fidelity of visual memory.

We observed a majority of visually responsive units that were recorded in ITC and HC showed RS, with only a few showed RE. Furthermore, RS units carried classifier predictive power and were matched to behavior. This is reminiscent of previous work from Sakon & Suzuki (2019), who reported that visual memory representations in HC were reflected as mixtures of RS in some units and RE in others, however, the units that reflected the most RS were able to discriminate between repeat and lure images whereas RE units were not. Of note is that the behavioral paradigm employed by Sakon & Suzuki centered on preferential looking (where monkeys innately prefer looking at novel images longer than repeated ones), thus combining a motivational component (curiosity) with familiarity. Together, these studies suggest that, like ITC, HC visual recognition memory representations are reflected as RS whereas RE may reflect other processes in these brain areas (see also (Kim, 2017)).

These results challenge the traditional view that the hippocampus uniquely shapes the fidelity of visual memory. They suggest that instead, the fidelity of visual memory follows from integrated computations across brain regions. Understanding memory failures (from diseases to typical errors) requires an understanding of visual memory that appreciates the specific and combined contributions of neocortical regions and the medial temporal lobe. This study helps build toward a holistic model of visual memory by demonstrating that computations traditionally thought to happen only in the hippocampus are reflected in visual cortex.

## Methods

### Subjects

Experiments were performed on two adult rhesus macaque monkeys (one female, *Macaca mulatta*) with implanted headposts and recording chambers. All procedures were performed in accordance with the guidelines of the University of Pennsylvania Institutional Animal Care and Use Committee.

### The modified mnemonic similarity task

All behavioral training and testing was performed using standard operant conditioning with juice as a reward. The monkeys’ eyes were tracked using high-accuracy infrared video eye tracking under head-stabilized conditions using Eyelink software and hardware (https://www.sr-research.com/). Image stimuli were presented on an 85 Hz refresh rate LCD monitor using customized software (http://mworks-project.org).

Trials were initiated by the monkey fixating on a red fixation square (.25°) at the center of a gray screen, within a square window of length 3.5°, followed by a 500ms delay before a (4°) full-color image appeared. The monkey maintained fixation on the image for 500ms before the red fixation square turned to a green go cue, and two targets were displayed 8° above and below the image. The monkey could then make a saccade to one of the two targets to indicate if that stimulus was novel (never before seen), or repeated (the exact same image, seen exactly once before). Images were never presented more than twice during the entire training and testing period of the experiment and fell into one of two categories. The first category was ’exact repeats’, where an image was shown once as novel and once as repeated. The second category were ‘lure pairs’ where, rather than being followed by a repeated presentation of an image, the image was followed by a ‘lure’ image showing the a novel image of the same object at titrated levels of similarity to the novel image. Because the lure images had never been seen before, they were rewarded as novel images. To keep the ratio of novel versus repeated images approximately equal, the lure images were repeated (these trials are not included in the analyses we report here).

Saccades made before the go cues resulted in an aborted trial (and were excluded from the analyses we report here). The association between the target (up vs. down) and the report (novel vs. repeated) was swapped between the two animals. For Monkey 1, the stimulus remained on the screen until a saccade away from the stimulus was detected. For Monkey 2, the stimulus disappeared at the end of the 500ms presentation.

In the experiment, the image repeats and lures were shown with an ’N-back’ (N=number of interleaving images) delay adjusted to approximately equate performance between the two animals, determined to be ∼24 images apart in monkey 1 and ∼8 images apart in monkey 2. The target N-back was chosen such that performance for exact repeats was near 100% and performance on lure image pairs was off ceiling (and modulated by dissimilarity). N-backs for individual trials were randomly selected from a uniform distribution of nearby N-backs (monkey 1: [23,24,25,26], monkey2: [7,8,9,10]). In addition, N-backs of [1,2,4] were shown throughout the experiment to serve both as fillers and as vigilance trials and a small fraction of trials were filled with other N-backs as fillers (e.g. 3-back). These trials were not included in the analyses reported here.

In total, a behavior session included 1,500 trials. Of these, 480 were associated with experimental lures (160 novel images, 160 visually similar lures and 160 lure repeats), 900 were exact repeat images shown at matched n-backs to the lures (450 novel images and 450 repeats), and 120 were vigilance lures (40 novel images, 40 visually similar lures and 40 lure repeats).

### Images

The images used in these experiments were gathered from five databases: the Ecoset image database (Mehrer et al., 2021), the Things image database (Hebart et al., 2019), the MemCat image database (Goetschalckx and Wagemans, 2019), the LaMem image database (Khosla et al., 2015), and the MST task (Kirwan et al., 2007; Bakker et al., 2008). The vast majority of the images used in these experiments were Ecoset images. Images were cropped and resized to a square of dimension 256*256 pixels. Image memorability (Isola et al., 2011; Khosla et al., 2015) has been shown to effect the response magnitude to observed images (Jaegle et al., 2019).

This poses a potential confound to interpreting magnitude-based memory signals. To control for this, images were run through an artificial network MemNet (Khosla et al., 2015) to yield a predicted memorability score for each stimulus. Because the memorability of an image is known to affect the magnitude of its neural response, and this project was not focused on disambiguating effects of image memorability, only high-memorability images with a MemNet score of .7-1 were considered for this project, and memorability distributions were matched between lure bins.

### Defining visual dissimilarity

There were two hurdles to overcome when defining visual dissimilarity. First, our experiment is designed to measure behavioral performance and its neural correlates relative to an ITC benchmark that we do not have direct access to before the experiment begins. Second, motivated by the robustness of single-trial familiarity behavior, our experiment focuses on the nature of familiarity representations in the brain following a single exposure, and the single-trial neural data we collect are insufficient to yield robust predictions of image dissimilarity for individual image pairs. To address these challenges, we turned to representational dissimilarity as defined by the dissimilarity of image representations in the brain-like artificial neural network CORnet_S-IT. CORnet_S-IT is a top-performing model evaluated for its ability to mimic the responses of ITC to arbitrary natural images in passively fixating monkeys, along with behavioral patterns of confusions in humans on a invariant object recognition task (Kubilius et al., 2019). Using methods described by Kriegeskorte et al., 2008, we created a representational n*n dissimilarity matrices, where n is the number of images within an object category, and each value is equal to 1 minus the Pearson’s correlation of two images’ representations in CORnet_S-IT. Of note, CORnet_S-IT model predictions are noiseless and thus the visual dissimilarity of an image to itself is 0. We expect dissimilarities of two non-paired novel images to be approximately 1. We verified this by computing the dissimilarity of each novel images to another (non-lure) novel image not confined to the same object category. This indeed resulted in a distribution approximated by 1.

From each image set, for each object category, a dissimilarity matrix was calculated by comparing each image within the category to each other image within that category. After using methods described above to generate a dissimilarity matrix for each image to all images within an object category, putative pairs were created using the two images with the lowest dissimilarity score. Pairs were sampled without replacement from the matrix until all images were paired, yielding more than 400,000 potential pairs at a range of dissimilarities. In the case of MST lure images, original image pairings were maintained, but dissimilarities were calculated like other image pairs. Selected pairs were spread evenly by dissimilarity across session while making sure that each object category was presented as lure pair no more than four times across an entire session.

After lure pairs were selected, extra images from unused pairs were used to populate each experimental session as exactly repeated images. These images were never shown in the same session as its potential pair.

### Neural recording

The activity of units in HC and ITC were recorded in two monkeys while they performed the modified mnemonic similarity task. Following on Sakon & Suzuki (2019), we targeted HC subregions CA3 and dentate gyrus. We do not have the resolution required to differentiate CA3 from dentate gyrus, and report both as HC.

In monkey 1, HC recordings were made in a single chamber in the left hemisphere, and ITC recordings were made across two chambers placed over left and right ITC. In monkey 2, all recordings in both HC and ITC were made using a single chamber over right hemisphere.

Chamber placement was guided by anatomical magnetic resonance images of the monkey’s brain. In monkey 1, chambers were centered 15.5 mm lateral of midline, and 19mm anterior to the ear canals, with a 2-deg angle perpendicular to midline oriented dorsomedial to ventrolateral. HC recordings were sampled from 13 locations spanning from 13-16mm anterior of ear canals, 16-21mm lateral of midline, and depths 8-12mm above the ventral surface of the temporal lobe in the left hemisphere. ITC recordings utilized 6 recording locations over both hemispheres spanning 15-18mm anterior to ear canals, 19-23mm lateral of midline, and of 2-5mm above the skull (following ventral surface of the temporal lobe).

In monkey 2, the chamber was centered at 10mm lateral of midline to the right, and 15mm anterior to the ear canals, with a 9-deg angle perpendicular to midline oriented dorsomedial to ventrolateral. Recordings in HC utilized 7 recording locations spanning from 11-13mm anterior of ear canals, 11-14mm lateral of midline, and at recording depths 8-12mm above the ventral surface of the temporal lobe in the right hemisphere. ITC recordings utilized 7 recording locations in the hemispheres spanning 9-12mm anterior to ear canals, 18-21mm lateral of midline, and depths of 2-5mm above the ventral surface of the temporal lobe.

Neural recordings were performed once the monkeys were fully trained (performance on novel/direct repeat images of above 80% accuracy, mnemonic performance modulated by n-back as reported previously in the lab (Meyer and Rust, 2018), and behavior was not disrupted by the introduction of lure images in practice sessions). For monkey 1, combined neural recordings and behavioral sessions occurred in two periods around 6 months apart. During each period, recordings took place 4-5 times a week and spanned 8 weeks. For monkey 2, combined neural recordings and behavioral sessions occurred in a single 8-week period.

Neural activity was sampled using a 24-channel U-probe (Plexon, Inc, Dallas, TX) with channels spaced vertically along the shaft of the probe 100μm apart. Continuous wideband neural signals were amplified, digitized at 30kHZ, and stored using the Grapevine Data Acquisition System (Ripple, Inc., Salt Lake City, UT). Neural data was sorted into spikes manually offline (Plexon offline Sorter). During sorting, between one and three candidate unit(s) were identified on each recording channel. Spike sorting was performed blind to any experimental conditions to avoid bias.

### Session Exclusion Criterion

A recording session was included in the analysis if: (1) The monkey completed a minimum session length of 1200 trials; (2) At least one unit passed a visual screen comparing the evoked mean firing rate for each unit across trials before and after stimulus onset (between the window of -300-0ms vs 50ms-350ms) using a t-test with a threshold of p<0.1. (3) Firing rates were stable over the course of a session through visual inspection; (4) The recording contained a minimum RS or RE magnitude of 2% for direct repeated images. In addition, (5) some sessions were eliminated to match memory between brain regions in the same monkey. This step was performed to roughly match on SCC predicted memory performance at the end of the image viewing window (200ms-500ms). To match memory information between the two data sets, the last 7 recorded sessions from monkey 1 ITC were not analyzed. For the HC populations: In monkey 1, 26 sessions were recorded and 9 were removed (4 and 5 due to criteria 1 and 4, respectively). In monkey 2, 17 sessions were recorded and 6 were removed (2, 2 and 2 due to criteria 1 and 2 4 respectively). For the ITC populations: In monkey 1, 18 sessions were recorded and 8 were removed (1,1, 2, and 4 due to criteria 1,2, 3, and 5 respectively). In monkey 2, 19 sessions were recorded and 5 were removed (1, 2 and 2 due to criteria 2, 3, and 4, respectively). The sample size was chosen to approximately match that of our previous work (Meyer and Rust, 2018; Mehrpour et al., 2021).

### Pseudopopulation generation

Each session was limited to 24 channels, and yielded too little data for an assessment of alignment between neural and behavioral performance. To overcome this challenge, we combined data across sessions into larger pseudopopulations using methodology introduced by Meyer and Rust (2018). When creating these pseudopopulations, we took trials at our target n-back (∼24 for monkey 1 and ∼8 For monkey 2) and aligned data across sessions, splitting the data into 4 conditions: 1) novel images in the “exact repeat” condition, 2) novel images in the “lure” condition, 3) repeated images in the “exact repeat” condition, and 4) lure images. Lure pseudopopulations were aligned by CORnet_S-IT dissimilarity. For novel/exact repeat images, the pseudopopulation images were randomly aligned, consistent withdissimilarity values of 0 .

Because different images were used in each session, aligning images in this way implicitly assumes that: 1) the total number of spikes evoked in a brain area is matched across different images, 2) the data recorded in any one session is a representative sample of those statistics, and 3) that the responses of the units recorded in different sessions are uncorrelated.

The number of trials in each pseudopopulation was limited to the length of the shortest session, meaning that each session that was longer than the shortest had trials ablated until it matched in size. In order to trim populations, images to be ablated were chosen randomly. This process yielded four pseudopopulations for each brain region for each monkey: two lure pseudopopulations of the responses to each of the lure image pairs, and two exact repeat pseudopopulations of the novel and repeated response to the image.

Finally, the number of units included in each population was matched for memory information using SCC classifier performance in the viewing window 200-500ms. This resulted in the following performances (mean ± s.e.): Monkey 1 — HC: 0.850 ± 0.00013, ITC: 0.865 ± 0.00011; Monkey 2 — HC: 0.758 ± 0.00018, ITC: 0.744 ± 0.00016. For Monkey 1, pseudopopulations consisted of 422 HC and 504 ITC units, each exposed to 213 repeated and 133 lure images. For Monkey 2, 340 HC and 595 ITC units were used, with 142 repeated and 80 lure images.

The analyses in Figure 6 used pseudopopulations created by combining data from both monkeys, meaning pseudopopulations were limited to the smaller of the two paired populations resulting in sizes. For the combined analysis, pseudopopulations included 747 HC units and 1,024 IT units, each exposed to 142 repeated and 80 lure images.

### d’

Unit d’ was calculated as the difference between a unit’s mean response to novel images (between 300ms-500ms) compared to the response to the same image when it was repeated, divided by the average standard deviation across both sets (Figure 3b-c).

### Total spike count and weighted linear decoders

Following methods described in (Mehrpour et al., 2021), pseudopopulation responses were decoded from HC and ITC using two linear decoding schemes. For both decoders, the population response was quantified as the vector 𝐱 containing spike counts on a given trial. To ensure that the decoder did not erroneously rely on visual selectivity, the decoder was trained on balanced pairs of novel/repeated trials in which monkeys viewed the same image (regardless of behavioral outcome or experimental condition).

*Cross-validated training and testing:*

We applied the same, iterative cross-validated procedure for each linear decoder. On each iteration of the resampling procedure, the responses for each unit were randomly shuffled within each experimental condition to ensure that artificial correlations (e.g. between units recorded within a sessions) were removed. For exact repeat images, this shuffling was done across the entire pseudopopulation. Lure images were separated into three bins based on ranked similarity before shuffling, resulting in three bins of different dissimilarity. Each iteration was cross validated by reserving a subset of direct repeat images equal to the number of images in each lure bin (46 for Monkey 1, 26 for Monkey 2, and 26 for the combined population; roughly 25% of the novel/repeat responses). The remaining direct repeat and novel trials were used to train one of the linear decoders to distinguish novel versus repeated images. A criterion was then generated serving as a threshold to classify the response to a given image as a novel or repeat. This threshold was then used to predict whether the withheld novel/repeat images, as well as the lure images, were novel or repeated. A neural prediction of the proportion of trials on which “repeated” would be reported was computed as the proportion of test trials in a given condition that took on a value less than a criterion. Finally, the predicted response pattern was rescaled by a rescaling parameter as a proxy for adjusting the population size to match performance. This rescaling factor was computed by adjusting direct novel and repeat performance to 100% accuracy. This rescaling factor was then applied to lure performance.

Specifically, all decoders in this study took the general form of linear discriminators. The class (novel/repeated) of a population response vector, 𝐱 was determined by the sign of:

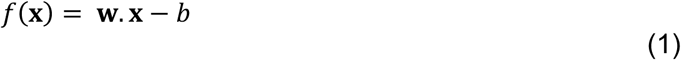

where **w** is an N-dimensional weight vector in the N-dimensional IT neural space (N is the number of units), and *b* is decision criterion, given by:

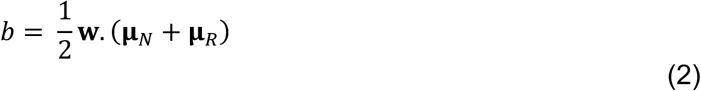

Here 𝛍*_N_* and 𝛍*_R_* are the mean population response vectors across novel and repeated images in the training set, respectively. A population response vector 𝐱 was classified as “novel” if 𝑓(𝐱) > 0, and “repeated” if 𝑓(𝐱) < 0.

The total spike count classifier (associated with RS) used a homogeneous weight vector (Figure 5-6):

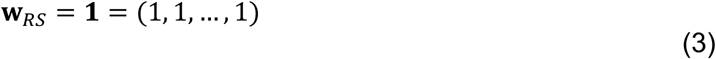

The weighted linear decoding vector (which could, in principle, combine a population of heterogeneous units in which some exhibit RS and others RE) was defined as (Figure 4):

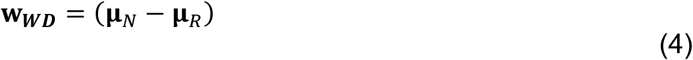

where 𝛍*_N_* and 𝛍*_R_* are the mean population response vectors across novel and familiar images in the training set, respectively. This is a simple form of a fisher linear discriminant that arises when the average covariance is a multiple of the identity, and is sometimes called a “prototype classifier”.

### Adding Ranked Units

Assessing the contributions of units to memory performance as a function of their ranked d-prime was analyzed by first generating the d-prime for each unit within a population based on the training data, and then adding units from a pool. This pool of units was tested for cross-validated performance as described above. For each run of the analysis, a ranking type (i.e. ranking by positive weights amplitude ‘High’; negative d-prime amplitude ‘Low’; or a random/shuffled ranking ‘Rand’) dictated which units were added. For each combination, classifier performance was gathered for each 10% of the total population (i.e. 10%, 20%, 30%… 100%) (Figure 4).

### Sharpening Metric

The goal of the sharpening metric was to quantify the degree of sharpening present in either a behavioral choice pattern or a prediction based on a neural classifier. This sharpening was quantified relative to a benchmark connecting the hit rate (repeated correct performance) to the false alarm rate (novel incorrect performance) by integrating the area of the curve with respect to this benchmark. To compare classifier sharpening and behavior, both classifier predictions and behavior were rescaled to 100% for repeats and novel images, and lure accuracy was rescaled proportionally. Specifically, this rescaling took classifier performance (specifically, average percent chose repeat) 𝐂 to all images and rescaled it using classifier performance to novel 𝐂*_N_* and repeat 𝐂*_R_* images as bounds to yield a rescaled percent chose repeat for all image dissimilarities 𝐂*_Rescaled_*.

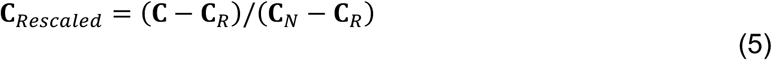

Next, the area below the curve relative to the linear benchmark was added, and area above the benchmark was subtracted. Under the assumption that lure performance is intermediate to novel and repeat performance, this value ranges from -1 to 1. Finally, to capture signal rather than noise, the result was weighted by the squared average performance of the classifier to novel and repeat images ranging from a weighting of 1 (at the highest observed classifier performance during the viewing period) to zero (when the classifier performance was at chance). The resulting sharpening metric ranged from -1 to 1, with positive scores representing sharpening, negative scores representing broadening, and 0 reflecting no change relative to the ITC benchmark (Figure 6).

## Financial interest or conflicts of interest

None.

## Acknowledgements

This work was supported by the National Science Foundation (award 2043255 to N.C.R), the National Eye Institute of the NIH (award R01EY020851 to N.C.R.) and the Simons Foundation (Simons Collaboration on the Global Brain award 543033 to N.C.R.).

## Supplemental Figures

**Supplemental Figure 1:**
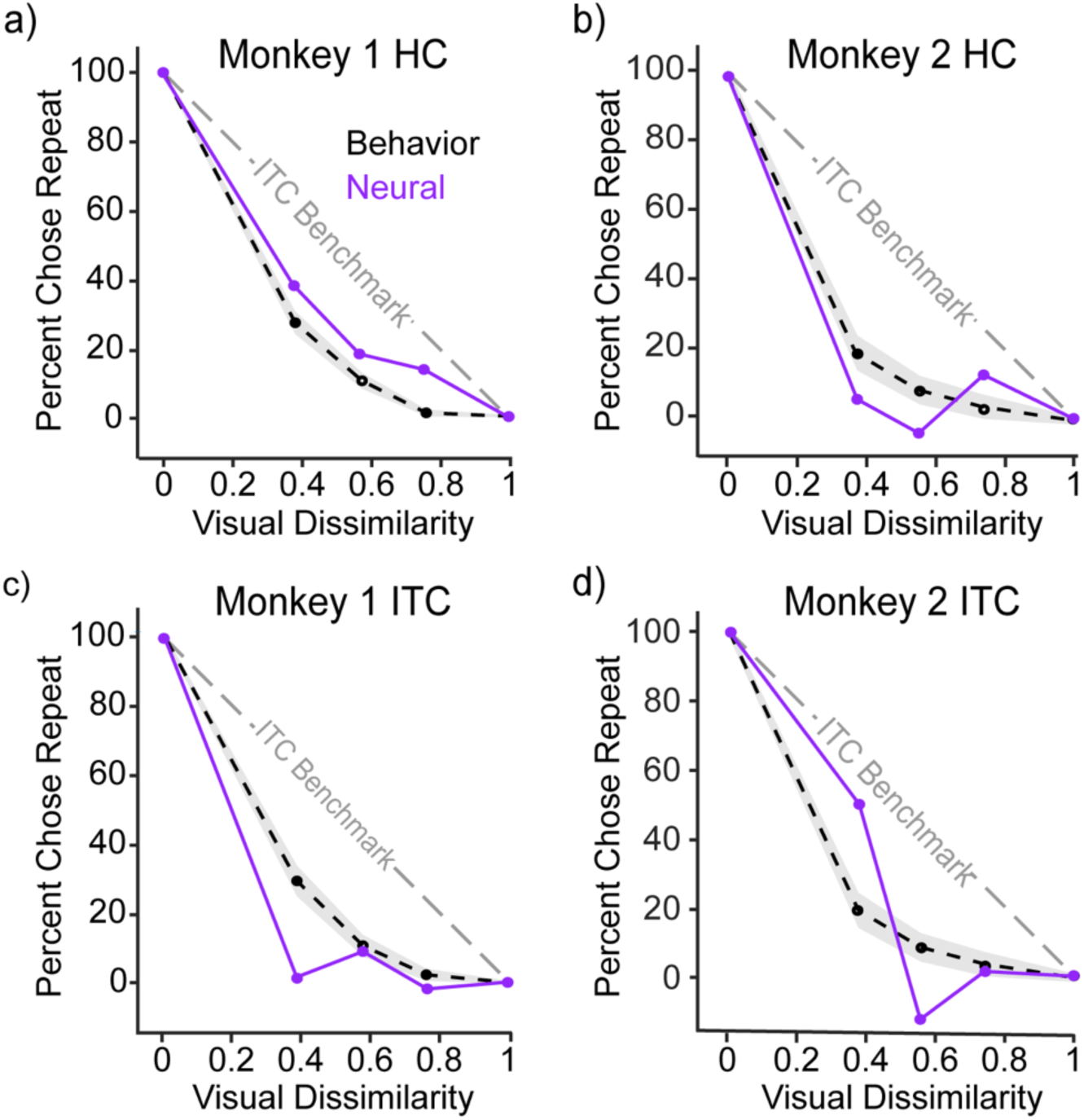
*Sharpening reflected in both HC and ITC is not dependent on extremes in firing rates.* Spike counts were z-scored for each unit across all images in order to determine if extremes in firing rates dominated classifier performance. We found that z-scoring had little effect on relative positions of distributions nor did it affect the match of classifiers to behavior. **(a-d)** Shown is the comparison of sharpening behavior (black, dashed) and cross-validated classifier performance (purple). Behavioral and classifier performance is plotted with the same conventions as Figure 5.

**Supplemental Figure 2.**
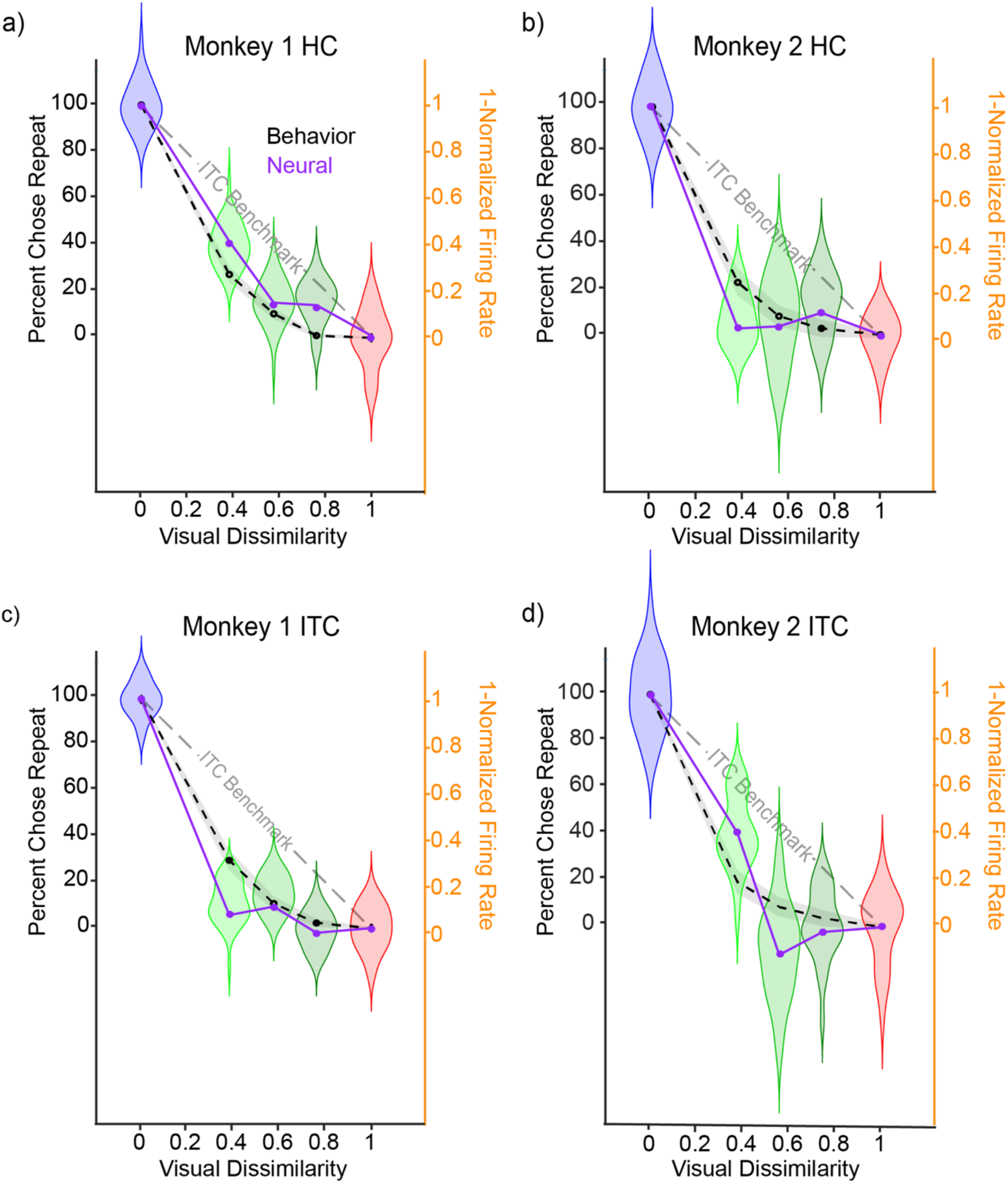
Firing rate Distributions match classifier performance HC and ITC. a-d) Behavior (black, dashed), cross-validated spike count classifier performance (purple), and firing rate vigor (violin plots) (blue/green/red for repeat/lure/novel respectively). Behavioral and classifier performance is plotted with the same conventions as Figure 5. Violin plots are 1-Normalized Firing Rate (righthand y-axis), reflecting smoothed distributions of firing rates for each dissimilarity condition. Note that in this rescaled space, values can exceed 1 and fall below 0, this rescaling allows for visualization of the distributions of firing rates compared to the classifier performance.

**Supplemental Figure 3:**
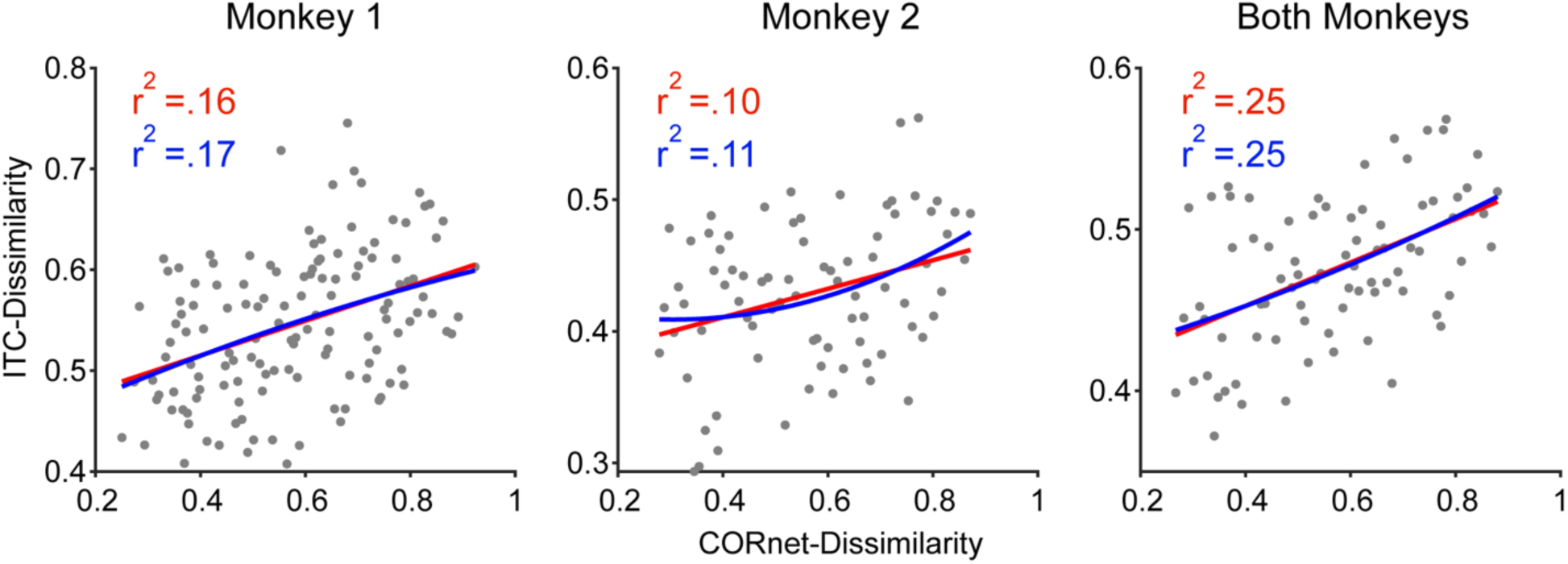
No evidence for a nonlinearity in CORnet_S-IT vs ITC dissimilarity. Shown are fits of ITC neural pairwise dissimilarities (1-pearson’s correlation of the pseudopopulation response to pseudoimages) as a function of CORnet_S-IT dissimilarities for each lure image within each pseudopopulation. Individual images are indicated as gray dots; Linear fits are indicated in red; quadratic fits are indicated in blue. Neural similarity was calculated using a window from 100ms-250ms, following the visual evoked response. Plots are shown for each monkey as well as pooled across the two monkeys. Of concern is a potential nonlinear relationship between these two quantities that might spuriously lead to the impression of sharpened visual memory neural representations and behavior; we find no evidence for this in the data.

